# VTA MC3R neurons control feeding in an activity and sex-dependent manner in mice

**DOI:** 10.1101/2021.03.09.434586

**Authors:** Anna I. Dunigan, David P. Olson, Aaron G. Roseberry

## Abstract

Increasing evidence indicates that the melanocortin and mesolimbic dopamine systems interact to regulate feeding and body weight. Because melanocortin-3 receptors (MC3R) are highly expressed in the ventral tegmental area (VTA), we tested whether VTA neurons expressing these receptors (VTA MC3R neurons) control feeding and body weight *in vivo*. We also tested whether there were sex differences in the ability of VTA MC3R neurons to control feeding, as MC3R −/− mice show sex-dependent alterations in reward feeding and dopamine levels, and there are clear sex differences in multiple dopamine-dependent behaviors and disorders. DREADD receptors were used to acutely activate and inhibit VTA MC3R neurons and changes in food intake and body weight were measured. Acutely altering the activity of VTA MC3R neurons decreased feeding in an activity- and sex-dependent manner, with acute activation decreasing feeding, but only in females, and acute inhibition decreasing feeding, but only in males. These differences did not appear to be due to sex differences in the number of VTA MC3R neurons, the ability of hM3Dq to activate VTA MC3R neurons, or the proportion of VTA MC3R neurons expressing tyrosine hydroxylase (TH). These studies demonstrate an important role for VTA MC3R neurons in the control of feeding and reveal important sex differences in behavior, whereby opposing changes in neuronal activity in male and female mice cause similar changes in behavior.

## Introduction

The melanocortin system is involved in the control of homeostatic (‘need-based’) feeding and is comprised of neurons in the arcuate nucleus of the hypothalamus that release the neuropeptides α-melanocyte-stimulating hormone (α-MSH; POMC neurons) or agouti-related peptide (AgRP neurons) as well as the central melanocortin-3 and melanocortin-4 receptors (MC3R and MC4R). Activation of melanocortin receptors or POMC neurons decreases food intake and body weight (Aponte, Atasoy, & Sternson, 2011; Fan, Boston, Kesterson, Hruby, & Cone, 1997; Pierroz et al., 2002), whereas AgRP neuron activation and MCR antagonism stimulates food intake and weight gain (Aponte et al., 2011; Graham, Shutter, Sarmiento, Sarosi, & Stark, 1997; Krashes et al., 2011; Ollmann et al., 1997; Rossi et al., 1998). Although MC4Rs have been extensively studied and play a clear role in the control of feeding, body weight and glucose homeostasis (Butler & Cone, 2002), much less is known about the exact role of MC3Rs. Both MTII, an MC3/4R agonist, and AgRP alter food intake in MC4R−/− mice (Marsh et al., 1999; Rowland, Schaub, Robertson, Andreasen, & Haskell-Luevano, 2010) demonstrating that MC3Rs can affect feeding. In addition, mice with mutated MC3Rs show altered fat mass, body weight, feeding efficiency, activity levels, fasting-induced refeeding, and food self-administration (Butler et al., 2000; Chen et al., 2000; Ghamari-Langroudi et al., 2018; Girardet et al., 2017; Lee et al., 2016; Mavrikaki et al., 2016; Renquist et al., 2012). Furthermore, cell-type specific manipulation of MC3R expression in dopamine or AgRP neurons also alters feeding and food reward (Ghamari-Langroudi et al., 2018; Girardet et al., 2017; Mavrikaki et al., 2016), although these data appear to be contradictory in some cases (i.e. see (Ghamari-Langroudi et al., 2018; Girardet et al., 2017)). Thus, there is evidence that MC3Rs regulate feeding and body weight, but overall, we still have a poor understanding of the exact role that MC3Rs play in the control of energy homeostasis.

In addition to controlling homeostatic feeding, the melanocortin system can also regulate reward-related (hedonic or ‘want-based’) feeding through its interactions with the mesolimbic dopamine system (Aaron G. Roseberry, Stuhrman, & Dunigan, 2015). For example, both POMC and AgRP neurons project to the VTA (Dietrich et al., 2012; Dunigan, Swanson, Olson, & Roseberry, 2020; King & Hentges, 2011), which is the center of the mesolimbic dopamine system, and MC3Rs are highly expressed in the VTA (Lippert, Ellacott, & Cone, 2014; Roselli-Rehfuss et al., 1993). MC3R −/− mice also show altered food self-administration (Mavrikaki et al., 2016) and altered sucrose preference and changes in dopamine turnover (Lippert et al., 2014), and our laboratory has shown that intra-VTA injections of melanocortin receptor agonists and antagonists altered both feeding and body weight, and sucrose intake in two-bottle choice tests and self-administration assays (A. G. Roseberry, 2013; Shanmugarajah, Dunigan, Frantz, & Roseberry, 2017; Yen & Roseberry, 2014). Interestingly the altered sucrose preference and dopamine turnover observed in the MC3R −/− mouse were sex dependent (Lippert et al., 2014), suggesting that the melanocortin-mesolimbic systems interaction may be sexually dimorphic.

Because MC3Rs are the predominant melanocortin receptor subtype in the VTA (Lippert et al., 2014), VTA neurons expressing this receptor may be a good candidate for a site of interaction between neural circuits controlling homeostatic and reward-based feeding. In these studies, we used transgenic mice expressing Cre recombinase in MC3R neurons (Pei et al., 2019) combined with DREADD receptor technology to test whether acute activation or inhibition of VTA MC3R neurons controls feeding and body weight and whether any observed changes in feeding and body weight were sex-dependent.

## Materials & Methods

### Reagents

Sterile bacteriostatic saline, ketamine, xylazine, and meloxicam were from Patterson Veterinary Supply, Inc. (Greeley, CO). Clozapine*-*N-oxide (CNO) was from Tocris, Inc. (Minneapolis, MN), and chlorpropamide was from BioVision, Inc. (Milpitas, CA). pAAV-hSyn-DIO-hM3Dq-mCherry and pAAV-hSyn-DIO-hM4Di-mCherry plasmids (Krashes et al., 2011) were direct gifts from Dr. Bryan Roth. pAAV-hSyn-DIO-mCherry (Addgene plasmid # 50459; RRID: Addgene_50459) was a gift from Bryan Roth and was obtained from Addgene. The pHelper and pAAV-RC (2/2) plasmids were a generous gift from Ralph DiLeone. All other reagents were from common commercial sources.

### Adeno-associated virus (AAV) preparation

AAVs were prepared using a triple transfection, helper-free method and were purified as previously described (Hommel et al., 2006). Briefly, HEK293 cells were transfected with equal amounts of pAAV (hM3Dq, hM4Di, or mCherry), pHelper, and pAAV-RC using standard calcium phosphate transfection procedures. Cells were collected ~80 hours post-transfection, resuspended in freezing buffer (150 mM NaCl, 50 mM Tris, pH 8.0), frozen and stored at −80°C until preparation. Cells then underwent 2 freeze-thaw cycles and were incubated with benzonase (50 U/ml final) at 37°C for 30 minutes. The lysate was added to a centrifuge tube containing a 15%, 25%, 40% and 60% iodixanol gradient and was spun at 184,000 x g (50,000 rpm in a Beckman Type 70Ti rotor) for 3 hours 20 minutes at 10°C. The 40% fraction was collected and exchanged with sterile 1X PBS using Amicon Ultra-15 Centrifugal Filter Unit Concentrators (100 kDalton; Millipore, Inc., Billerica, MA). Viral titers were calculated using the AAV pro Titration Kit (Clontech, Inc.) per the manufacturer’s instructions. The final purified viral particles were aliquoted and stored at −80°C, except during use, when they were stored at 4°C.

### Animals

Male and female transgenic mice (12-15 weeks old, 18-22 g females and 25-30 g males) expressing Cre recombinase in MC3R neurons (‘MC3R-Cre mice’) or expressing EYFP in MC3R neurons on a mixed C57/129 background were used in all experiments. MC3R-Cre mice were generously provided by David Olson (University of Michigan, Ann Arbor), and have been previously characterized and validated (Pei et al., 2019; West, Lu, Olson, & Roseberry, 2019). Mice expressing EYFP in MC3R neurons were generated by crossing MC3R-Cre mice with Ai3 transgenic mice expressing Cre inducible EYFP (The Jackson Laboratory, stock # 007903). Mice were housed in ventilated polycarbonate Animal Care System cages in a temperature- and humidity-controlled room under a 12:12 light/dark cycle (lights on at 6:00 or 7:00 am) with *ad libitum* food and water throughout the experiment. Mice were group housed until 2 weeks prior to surgery and were housed individually thereafter. All protocols and procedures were approved by the Institutional Animal Care and Use Committee at Georgia State University and conformed to the NIH *Guide for the Care and Use of Laboratory Animals*.

### Stereotaxic surgery

AAV’s were injected into the VTA of MC3R-Cre mice using standard flat-skull stereotaxic techniques. 7-10 week old mice were anesthetized with isoflurane (1–5%) and placed in a stereotaxic apparatus (David Kopf Instruments, Tujunga, CA). The VTA was targeted using the following coordinates (relative to bregma): A/P-3.3, M/L+/-1.32, DV-4.55 and 4.45 from the skull surface, at a 12° angle to the midline. 150 nL of AAV solution was injected bilaterally at each of the two depths (300 nL total/side) at a speed of 100 nl/min using a Nanoliter 2010 microinjector (World Precision Instruments, Sarasota, FL) fitted with glass pipettes with ~30-60 μm diameter tips. The pipettes were left in the brain for 3 minutes following the first injection and 5 minutes following the second injection to allow for AAV diffusion. For pain management, mice received meloxicam (1 mg/kg) at the onset of the surgery and again 24 hours post-surgery. The mice were given 5 weeks for recovery to allow for full DREADD expression prior to the onset of testing.

### Food intake and body weight measurement

Standard laboratory chow (Purina rodent diet #5001, PMI Nutrition International; 3.36 kcal/g) was used in all experiments. Mice were acclimated to eating out of a 10 cm petri dish placed at the bottom of a cage for 1 week prior to the onset of the experiments. For measurement of food intake, mice were given a set amount of food which was then weighed at specific time points to determine the amount of food eaten. For 24-hour food intake measurements, the bedding was sifted to account for spillage whereas spillage was not accounted for during the acute time point measurements due to time constraints and to reduce interference during the experiment. For measurement of body weight, mice were placed in a small box, weighed and then kept in the box for a short duration (<1 min) while the food was weighed before being returned to their cage.

### Acute CNO treatment

Mice were separated into body weight-matched groups prior to surgery and were then randomly injected with one of the three AAVs into the VTA. Following a 5-week recovery period baseline body weight and food intake were measured daily 1 hour before dark onset for 5 days before the first injection and on each intervening non-test day. On test days, body weight and food intake were measured 60 minutes prior to dark onset and all food was removed from the cage. Mice then received an intraperitoneal (IP) injection of CNO (1 mg/kg) or vehicle starting 30 minutes prior to dark onset, and food was re-introduced into the cage 3 minutes before dark onset. Food intake was measured at 1, 2, 3, 4, and 24 hours post-injection and body weight was measured 24 hours post-injection. The animals were allowed 2 days to recover and the second treatment was administered. Injections were performed in a counterbalanced manner to control for injection order with half of the mice receiving CNO injections on the first test day and the other half receiving CNO on the second test day. After the termination of the study the mice were euthanized via transcardiac perfusion and brains were collected for *post-hoc* analysis of DREADD receptor expression. CNO was initially dissolved in DMSO, diluted with sterile saline (to 10% DMSO), aliquoted and stored at −20°C until use. Upon use, CNO was diluted to 0.1 mg/mL with sterile saline (1% DMSO final) and used for the injections. The same DMSO/saline mixture (1% DMSO) was used as the vehicle control for all injections. IP injections were done in a volume of 10 μL/g body weight and the injection sides were alternated between each injection. The researcher performing all aspects of the study was blind to the animals’ treatment group until all of the data had been collected and DREADD/mCherry expression was confirmed. All mice excluded from the study (see below) were excluded prior to unblinding of the data.

### Histological confirmation of DREADD expression

At the end of all experiments, mice were deeply anesthetized with ketamine/xylazine (93/7 mg/kg) and transcardially perfused with ice cold phosphate buffered saline (PBS) followed by 4% paraformaldehyde. The brains were dissected, post-fixed with 4% paraformaldehyde at 4°C overnight, washed with 1X PBS and incubated in 30% sucrose (in 1X PBS) for 2-3 days until fully saturated with sucrose. The brains were then flash frozen in ethanol/dry ice cooled isopentane and stored at −80°C until sectioning. 40 μm thick coronal sections were collected at 200 μm intervals through the entire VTA on a cryostat, mounted on glass slides and coverslipped using mounting media containing 10% 1,4-diazabicyclo[2.2.2]octane (DABCO). Images were collected at 20x on an Olympus BX41 fluorescent microscope equipped with an Olympus DP73 camera. The images were collected in a grid to visualize the entire VTA and were stitched together post-acquisition using the ImageJ *Stitching* plugin (Preibisch, Saalfeld, & Tomancak, 2009). The number of mCherry positive neurons in the VTA and outside of the VTA were manually quantified using ImageJ *Cell counter* plugin (Schindelin et al., 2012).

### Excitatory DREADD (hm3Dq)-mediated activation of c-fos

Male and female MC3R-Cre mice were injected with AAV-hSyn-DIO-hM3Dq-mCherry or AAV-hSyn-DIO-mCherry into the VTA for analysis of CNO-induced *c-fos* expression. Four to five weeks later, mice received an IP injection of CNO (1mg/kg) or vehicle and were transcardially perfused 2 hours later. Mice used for sex difference analysis received injections of CNO or vehicle near dark onset (at the same time as the experimental injections for the measurement of food intake) and were perfused 2 hours later. The brains were dissected, processed, and the sections were collected as described above. c-fos was labeled using standard immunohistochemical (IHC) techniques. In short, brain sections were blocked for 6 hours at room temperature in blocking buffer (5% normal goat serum, 0.2% Triton X-100, 0.1% bovine serum albumin in 1X PBS), washed in 1X PBS for 5 minutes, and were incubated with rabbit anti-c-fos antibodies (Cat. # ABE457Millipore, Inc., Billerica, MA) diluted 1:1000 in antibody incubation buffer (0.2% Triton X-100, 1% bovine serum albumin in 1X PBS) overnight at 4°C. Sections were washed with 1X PBS 3 times for 5 minutes each and were incubated with Alexa Fluor 488 conjugated goat anti-rabbit antibodies (Catalog # A-11008, Invitrogen, Carlsbad, CA) diluted 1:1000 in antibody incubation buffer for 4 hours at room temperature. Sections were then washed with 1X PBS 3 times for 5 minutes each, mounted on glass slides and coverslipped with mounting media containing DABCO. Images of VTA sections containing mCherry and Alexa Fluor 488 (c-fos) signals were acquired using a 20x objective with 0.7X zoom on a confocal microscope (LSM 720; Carl Zeiss, Oberkochen, Germany). The numbers of neurons expressing mCherry, c-fos, and those co-expressing mCherry and c-fos were manually quantified using the ImageJ *Cell Counter* plugin (Schindelin et al., 2012).

### MC3R expression analysis

Brains of male and female mice expressing EYFP in MC3R neurons were collected and processed as describe above and neurons expressing TH were labeled using standard immunohistochemical (IHC) techniques (as described above for c-fos) using mouse anti-TH antibodies (Catalog # MAB318, Millipore, Inc., Billerica, MA) diluted 1:1500 followed by Alexa Fluor 594 conjugated goat anti-mouse antibodies (Catalog # A-11032, Invotrogen, Carlsbad, CA) diluted 1:1000 in antibody incubation buffer. Images of the entire VTA (−2.92 mm to −4.16 mm from bregma) were acquired using a 20x objective with 0.7X zoom on a confocal microscope (LSM 720; Carl Zeiss, Oberkochen, Germany) and were stitched for each VTA section using XuvTools v.1.8.0 (Emmenlauer et al., 2009). The numbers of EYFP positive (MC3R) neurons, Alexa Fluor 594 positive (TH) neurons, and co-expressing neurons were quantified for each section using IMARIS v.9.5.1 (Bitplane). Total cell counts, cell counts per each VTA section, and cell counts for rostral (−2.92mm to −3.28mm from bregma), middle (−3.40 to −3.80mm from bregma) and caudal (−3.88 to −4.16mm from bregma) VTA were compared between the two sexes.

### Data analysis, statistics, and experimental design

All data are presented as means ± SEM with individual data points included in each graph. Data were graphed using IgorPro (Wavemetrics, Inc., Lake Oswego, OR), and statistical analyses were performed using IBM SPSS Statistics for Windows, Version 25.0 (Armonk, NY). Cumulative food intake was analyzed using a general linear model with repeated measures, with group (hM3Dq, hM4Di, mCherry control), treatment (vehicle *vs.* CNO), time, and sex (male *vs.* female), as independent variables. Percent decreases in food intake in males *vs.* females treated with vehicle *vs.* CNO were analyzed using general linear model with repeated measures with time and sex (male *vs.* female) as independent variables. All ANOVAs were followed by *post-hoc* tests corrected for multiple comparisons: Sidak. One-way ANOVA analyses were used to make sex comparisons of DREADD-mediated *c-fos* expression and MC3R expression. A significance level of p<0.05 was set *a priori* for all analyses.

For the behavioral experiments, mice were included in the data analyses only if the following criteria were met: DREADDs were bilaterally expressed in the VTA, the mouse did not have a clearly visible brain lesion at the injection site, and off-target (e.g. outside the VTA) labeling was <15% of the total number of cells labelled. Some individual mice had poor injections with only a few labeled neurons. To ensure that we only included mice with sufficient labeling to produce a behavioral response, we compared the total number of labeled neurons identified in the VTA in an individual mouse to the mean labeling for the group and excluded mice in which the labeling was 1 standard deviation below the group mean. Using these criteria resulted in exclusion of 47 mice total (7 control, 20 Gq, 20 Gi). To calculate the means for DREADD-expressing neurons, Gi and Gq groups were combined to obtain a common mean since these groups showed similar expression levels. The control group mean was standalone because the mCherry expression in this group was much higher than in the other two groups. These selection criteria were set *a priori*, and all mice were excluded prior to data unblinding and analysis.

The experiments were repeated in 7 independent cohorts of mice, with the size of the cohorts ranging from 9 to 16 mice per cohort. Information on all of the cohorts of mice tested in these studies, including those that were excluded, is provided in Table 1. This information is included for full transparency and reproducibility.

**Table 1:**
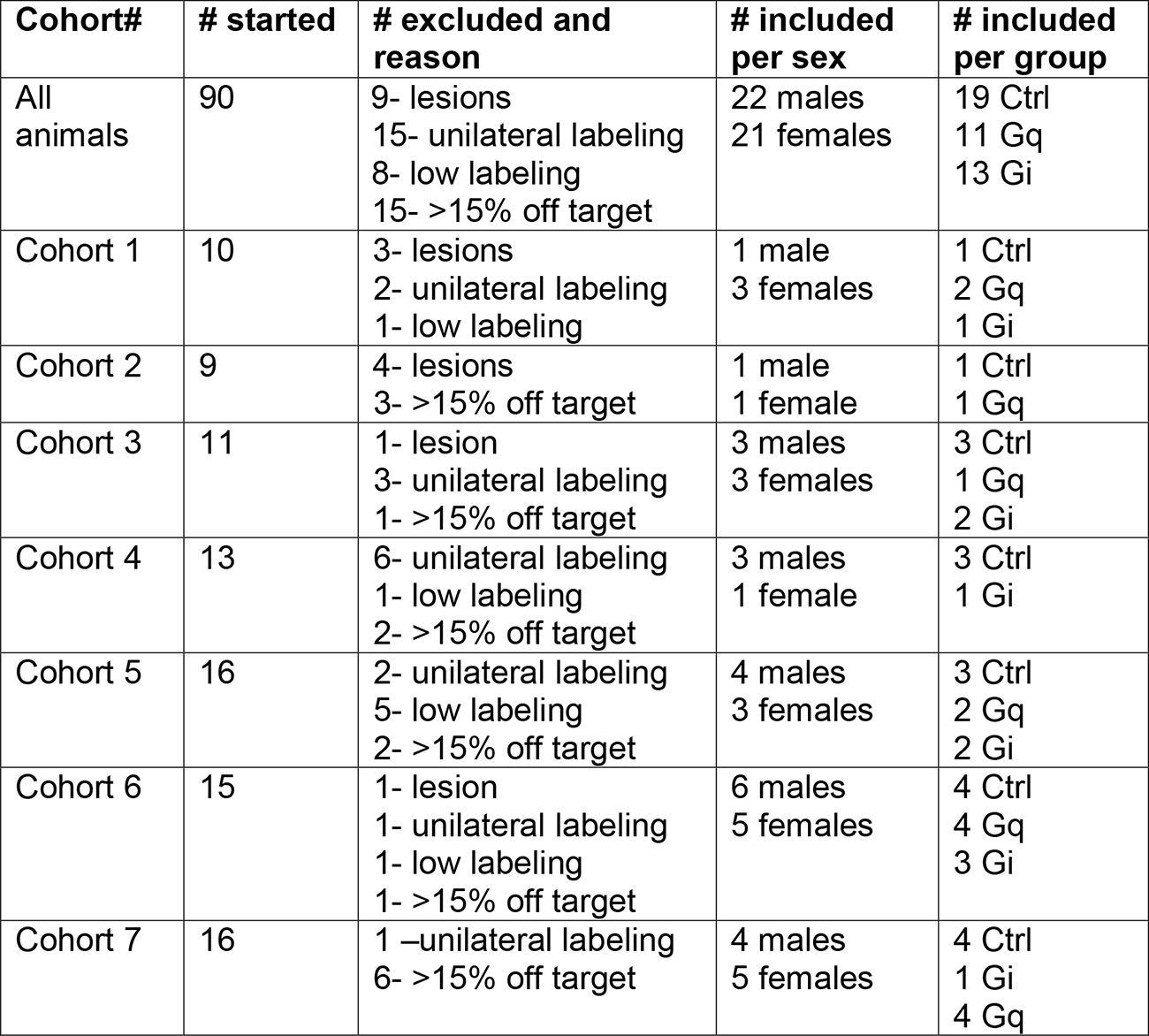
Detailed description of experimental animals used and excluded in the feeding experiment.

## Results

In these studies, we tested whether acute activation or inhibition of VTA MC3R neurons affected feeding and body weight and whether any observed effects were sex-dependent. AAVs expressing the excitatory DREADD receptor, hM3Dq-mCherry, the inhibitory DREADD receptor, hM4Di-mCherry, or the control, mCherry alone, in a Cre-dependent manner were bilaterally injected into the VTA of MC3R-Cre mice (Figure 1A, B) (Pei et al., 2019; West et al., 2019). CNO (1 mg/kg ip) was then used to activate or inhibit VTA MC3R neurons and changes in feeding and body weight were measured. For all experiments, DREADD-mCherry expression was confirmed to be restricted to the VTA for each mouse at the end of the experiment, and mice were excluded from analysis if the injections were unilateral, there were lesions at the injection site, or if there was significant mCherry expression outside the VTA (Table 1). Representative images showing VTA expression of mCherry for each group are shown in Figure 1C-E, and the mCherry expression pattern was consistent with that shown previously for MC3R expression in the VTA (Lippert et al., 2014; West et al., 2019)

**Figure 1:**
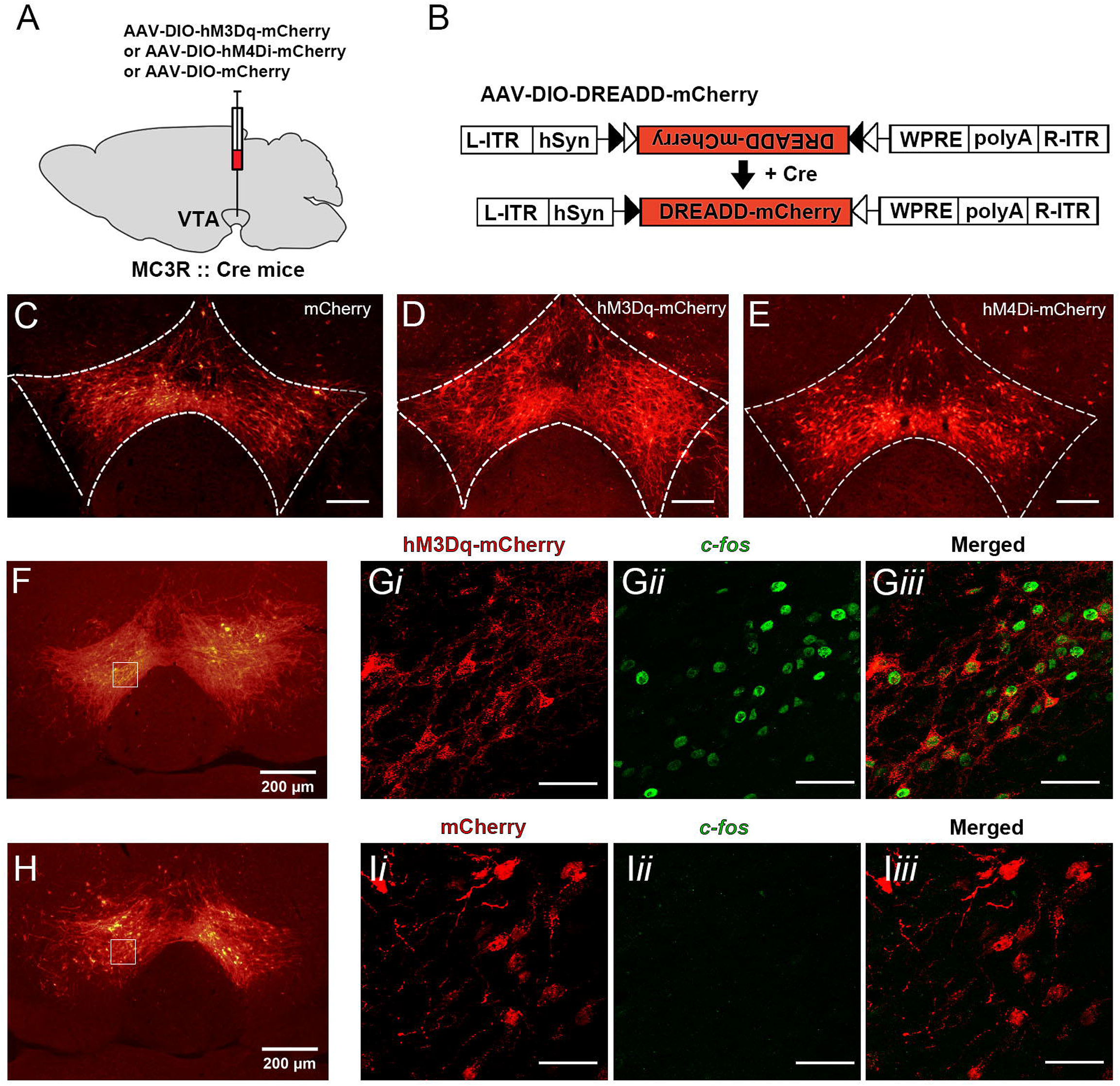
Confirmation of DREADD receptor expression and DREADD-mediated activation of VTA MC3R neurons in MC3R-cre mice. **A.** Graphical representation of AAV-DREADD delivery into the VTA. **B.** General experimental design for AAV-DIO-hM3Dq-mCherry, AAV-DIO-hM4Di-mCherry, and AAV-DIO-mCherry expression. **C-E.** Representative VTA sections showing mCherry control (C), hM3Dq-mCherry (D), and hM4Di-mCherry (E) expression in the VTA of MC3R-cre mice. **F-I.** c-fos immunoreactivity two hours after CNO injection (1 mg/kg, i.p.) in VTA neurons expressing hM3Dq-mCherry (F, G) or mCherry control (H, I). Boxes in F and H delineate anatomical location of high magnification images in G and I, respectively. Scale bars: 100 μm (C-E), 50 μm (G, I). L-ITR, left-inverted terminal repeat; hSyn, human synapsin promoter; R-ITR, right-inverted terminal repeat; WPRE, woodchuck hepatitis post-transcriptional regulatory element.

Although the ability of DREADD receptors to excite and inhibit VTA neurons has been previously established through electrophysiology and IHC (Runegaard et al., 2018; Sandhu et al., 2018; S. Wang, Tan, Zhang, & Luo, 2013), we confirmed hM3Dq’s ability to excite VTA MC3R neurons by examining changes in *c-fos* expression following CNO administration. CNO greatly increased c-fos in the VTA neurons expressing hM3Dq-mCherry (Figure 1F-G) compared to the VTA neurons from the control mice expressing mCherry (Figure 1H-I). Consistent with a previous report (S. Wang et al., 2013), we also observed an increase in c-fos in the VTA neurons lacking hM3Dq-mCherry, suggesting excitation through local connectivity or circuit network activity (Figure 1G).

We then tested whether acute activation or inhibition of VTA MC3R neurons with CNO (1 mg/kg) at the onset of the dark phase affected home cage chow consumption or body weight. Although there were small reductions in feeding and body weight when all mice were analyzed together (independent of sex) for both activation and inhibition of VTA MC3R neurons, there were no significant differences observed (data not shown). Addition of sex as a variable in the analysis revealed significant sex-specific effects for both activation and inhibition, however, as shown by significant treatment*group*sex and time*treatment*group*sex interactions. *Post-hoc* analyses demonstrated that activation of VTA MC3R neurons significantly decreased intake in female mice at all time points (Figure 2D) but did not affect the intake in male mice (Figure 2C). In contrast, inhibition of VTA MC3R neurons significantly decreased food intake in male mice but had no effect in females (Figure 2E-F). As expected, there were no effects of CNO in control mice of either sex (Figure 2A-B).

**Figure 2:**
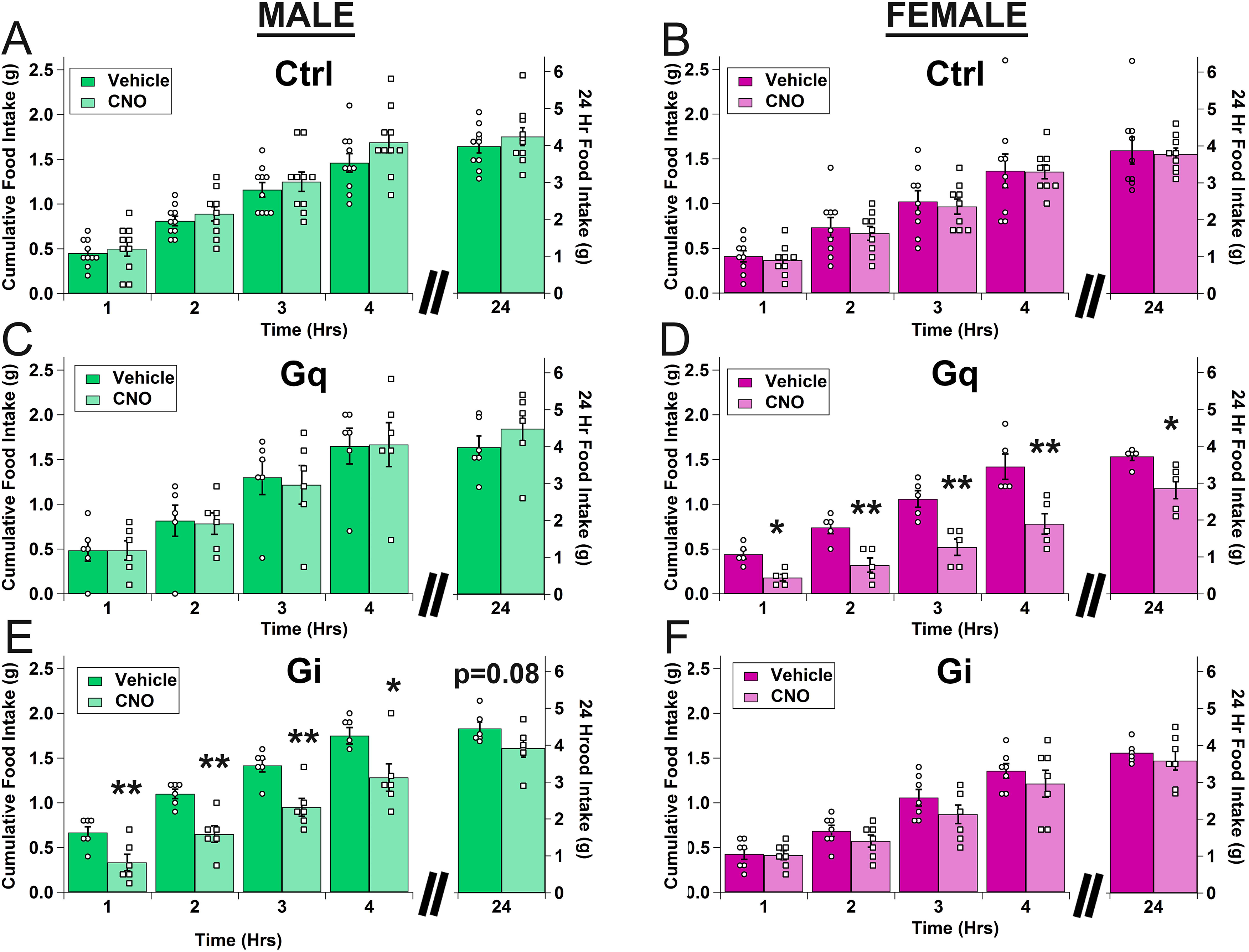
Effects of acute activation and inhibition of VTA MC3R neurons on food intake in males and females. **A-F.** Mean cumulative food intake for male (A, C, E) and female (B, D, F) mice (green = male, magenta = female). **A, B**. Control (mCherry). **C D**. VTA MC3R neuron activation with hM3Dq (‘Gq’). **E, F**. VTA MC3R neuron inhibition with hM4Di (‘Gi’). Control: n=19 (10 males, 9 females), Gq: n=11 (6 males, 5 females), Gi: n=13 (6 males, 7 females). Open circles are individual datapoints for vehicle and open squares are individual datapoints for CNO.* p<0.05 *vs.* vehicle, **p<0.005 *vs.* vehicle.

To further examine the sex differences observed in the responses to VTA MC3R neuron activation and inhibition we compared the percent decrease in food intake produced by CNO (relative to vehicle) in males *vs*. females for each DREADD group. Although we did not observe a significant main effect for activation of VTA MC3R neurons, *post hoc* tests planned *a priori* revealed that activation of VTA MC3R neurons decreased feeding to a greater extent in females compared to males (Figure 3B). For the inhibition of VTA MC3R neurons a significant time*sex interaction was observed, and *post hoc* analyses revealed a trend towards a greater CNO-induced decrease in food intake in males at 1- and 2-hour time points but these values did not reach statistical significance likely due to the high variability in the response in females (Figure 3C). As expected, there were no sex differences in the percent decrease in food intake produced by CNO in the control groups (Figure 3A). Thus, these data clearly demonstrate that VTA MC3R neurons acutely control feeding in a sex- and activity-dependent manner.

**Figure 3:**
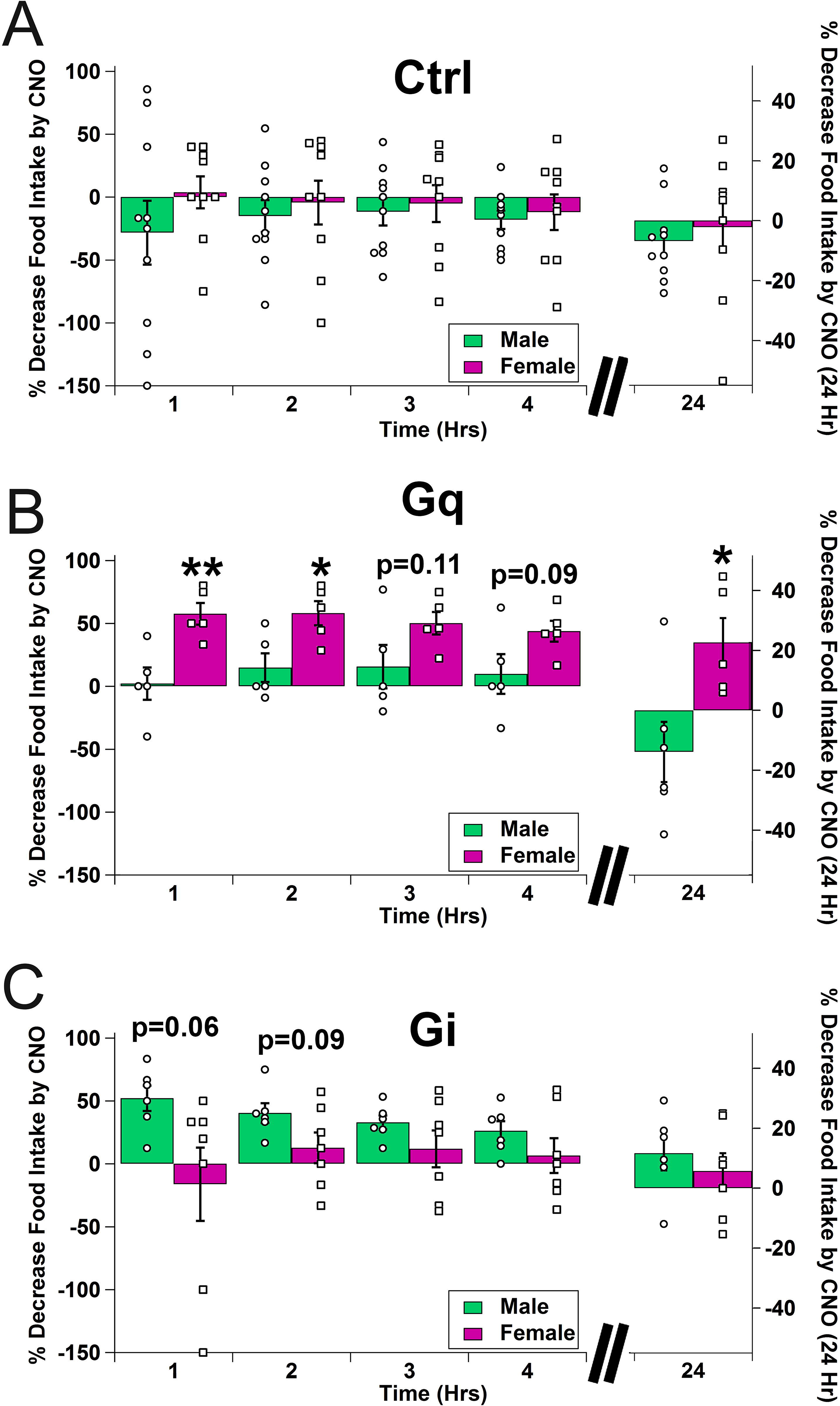
Comparison of the decrease in food intake produced by CNO between males and females. **A-C**. Percent decrease in food intake produced by acute CNO injection relative to vehicle in males (green) and females (magenta) expressing control (**A**), Gq **(B**), or Gi (**C**) DREADD in VTA MC3R neurons. Control: n=19 (10 males, 9 females), Gq: n=11 (6 males, 5 females), Gi: n=13 (6 males, 7 females). Open circles are individual datapoints for males and open squares are individual datapoints for females. * p<0.05 *vs.* vehicle, **p<0.005 *vs.* male

We next conducted a series of experiments to try to identify the potential cause of the sex differences observed in our behavioral experiment. To determine if the observed differences could be due to sex differences in the ability of hM3Dq to activate VTA MC3R neurons, we compared the number of VTA neurons co-expressing hM3Dq and c-fos following CNO or vehicle injection in male (Figure 4A) and female (Figure 4C) mice expressing hM3Dq-mCherry in VTA MC3R neurons. As we observed in the control experiments in Figure 1, CNO increased c-fos in the VTA neurons expressing hM3Dq-mCherry (Figure 4A-D) compared to vehicle injection in both males and females, but there was no difference in the number of VTA MC3R neurons co-expressing c-fos between sexes (Figure 4E). Thus, it appears that these effects were not due to differences in the ability of DREADD receptors to activate VTA MC3R neurons between sexes.

**Figure 4:**
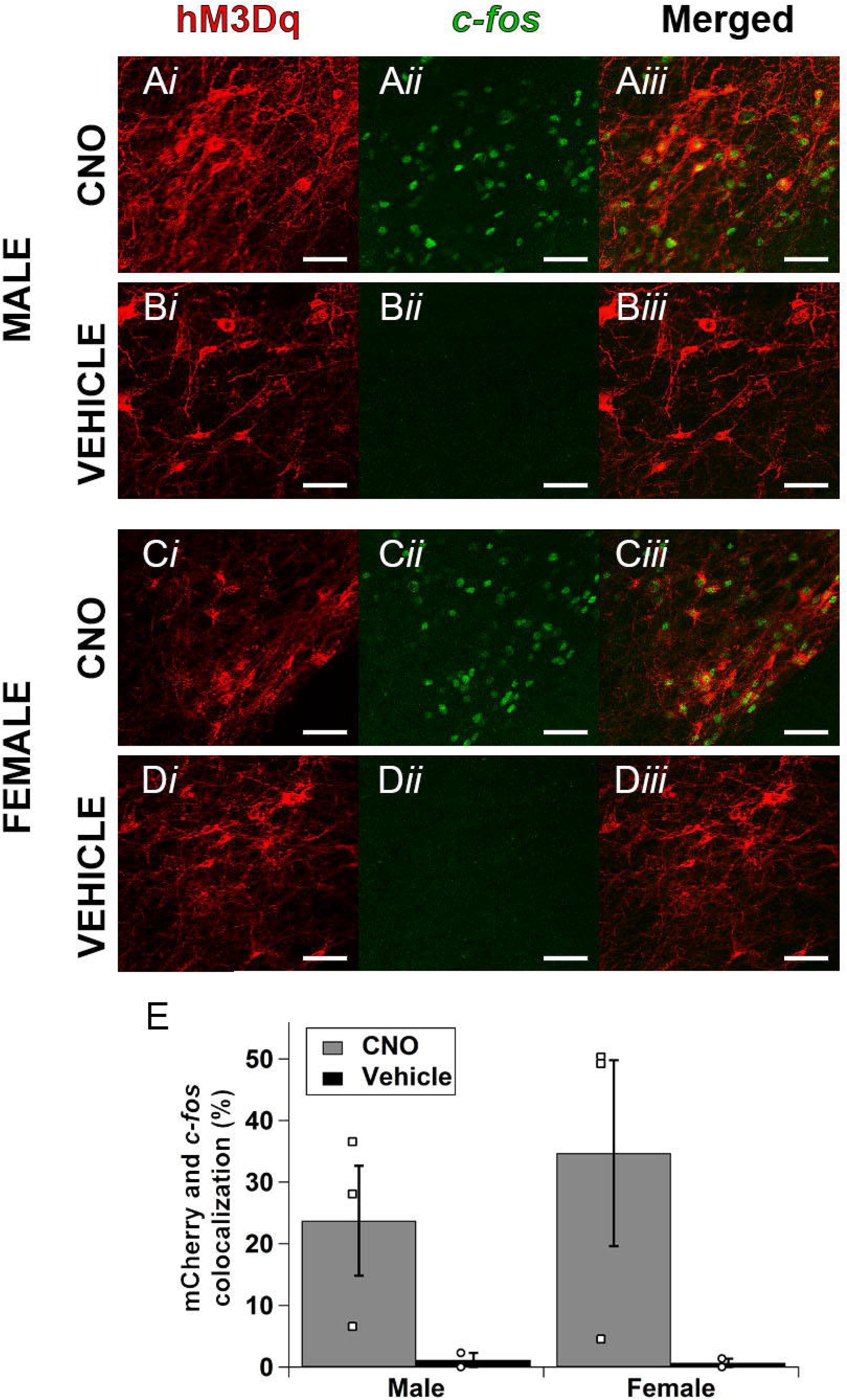
Comparison of CNO-mediated activation of *c-fos* in males and females. **A-D**. Sample images from male (A, B) and female (C, D) mice treated with CNO (A, C) or vehicle (B, D). **E**. Mean mCherry and c-fos colocalization in male and female mice treated with CNO (gray) or vehicle (blue) expressed as percent of total mCherry-expressing neurons. Male and female CNO: n=3, male and female vehicle: n=2. Open squares are individual datapoints for CNO and open circles are individual data points for vehicle. Scale bars are 50 μm.

We next tested whether the number of VTA MC3R neurons that are dopamine neurons differed between sexes by analyzing the expression of TH in VTA MC3R neurons. Sample VTA sections collected from male and female mice expressing EYFP in MC3R neurons (green) and immunolabeled for TH (red) are shown in Figure 5. MC3R-expressing neurons were found along the rostro-caudal extent of the VTA but were more concentrated in the middle VTA (−3.40mm to −3.64mm from bregma, Figure 5D) in both sexes. There were no differences in the total number of neurons in the VTA that expressed EYFP (data not shown) or the number of MC3R neurons at different locations along the rostro-caudal extent of the VTA between sexes (Figure 5D). There were also no differences in the total number of TH+ neurons or their rostro-caudal distribution between sexes (data not shown). Importantly, the number of VTA MC3R neurons co-expressing TH (Figure 5B) and the number of TH-positive neurons co-expressing MC3Rs (Figure 5C) were not different between the two sexes, although there was some variability between mice, especially in the middle VTA. Among the MC3R positive population, 37 ± 2, 64 ± 3%, and 52 ± 4% co-expressed TH in rostral, middle, and caudal sections, respectively, in males and females combined (Figure 5B), whereas 16 ± 2%, 28 ± 3%, and 25 ± 3% of TH positive neurons co-expressed MC3R in rostral, middle, and caudal sections (Figure 5C). Thus, it appears that the sex differences in the effects of activation and inhibition of VTA MC3R neurons on feeding were not due to differences in the total number VTA MC3R neurons, the total number of TH neurons or the proportion of MC3R neurons co-expressing TH.

**Figure 5:**
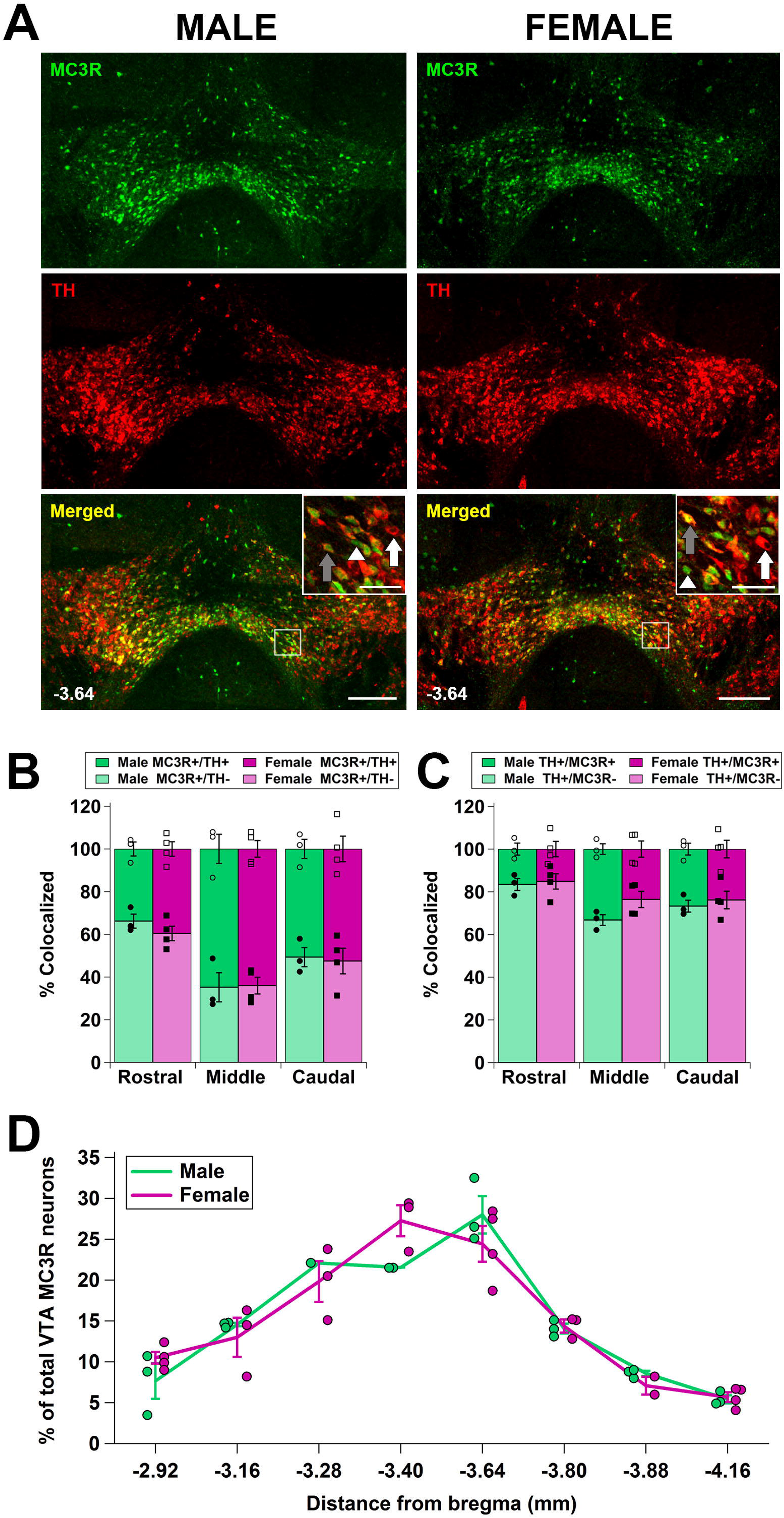
Comparison of neurons co-expressing MC3Rs and TH in males and females. **A**. Sample images from male and female mice immunostained for TH (red) and expressing EYFP in MC3R neurons (green). Insets show higher magnification of the regions in the box and show examples of MC3R+/TH+ (gray arrow), MC3R+/TH-(white arrowhead), and MC3R-/TH+ (white arrow) neurons. **B**. Percentage of MC3R neurons that are positive (dark bar) or negative (light bar) for TH in rostral (−2.92mm to −3.28mm from bregma), middle (−3.40gmm to −3.80mm from bregma), and caudal (−3.88mm to − 4.16mm from bregma) VTA section of male (green) and female (magenta) mice. **C**. Percentage of TH positive neurons that do (dark bar) or do not (light bar) contain MC3R-EYFP in rostral, middle, and caudal VTA sections from male (green) and female (magenta) mice. **D**. Rostro-caudal distribution of MC3R-expressing VTA neurons in males (green) and females (magenta). Male: n=3, female: n=4. Open circles and squares (B, C) and colored circles (D) are individual data points. Scale bars: 200 μm (VTA sections), 50 μm (insets).

## Discussion

In these studies, we tested the effects of acute activation and inhibition of VTA MC3R neurons on food intake and body weight and analyzed whether there were sex differences in the responses to changes in VTA MC3R neuron activity. There were interesting and divergent responses to acute VTA MC3R neuron activation and inhibition, with activation decreasing feeding only in females, and inhibition decreasing feeding only in males (Figure 2) demonstrating that VTA MC3R neurons acutely control feeding in an activity- and sex-dependent manner. We also showed that these sex differences in feeding were not due to sex differences in the ability of Gq DREADD receptors to activate VTA MC3R neurons, the total number of VTA MC3R neurons, or the percentage of VTA MC3R neurons co-expressing TH as a marker for dopamine neurons.

The most interesting result of these studies was the robust sex difference in the response to acute activation and inhibition of VTA MC3R neurons (Figure 2, 3). To our knowledge, this is the first demonstration that opposing changes in the activity of a specific neuronal population in males and females can cause the same behavioral response (decreased feeding). Although there is sexual dimorphism in the regulation of feeding and energy homeostasis (Asarian & Geary, 2013; Shi, Seeley, & Clegg, 2009) and sex- and estrous cycle-dependent variations in the melanocortin (Hubbard et al., 2019; Stincic, Rønnekleiv, & Kelly, 2018) and mesolimbic dopamine systems (Becker, 2016; Calipari et al., 2017; Morissette & Di Paolo, 1993), sex differences in behavior, including the regulation of feeding and body weight, typically manifest as a difference in the magnitude of the response between sexes rather than responses in opposite directions. For example, deletion of *Sirt 1* (Ramadori et al., 2010) or *STAT3* (Xu, Ste-Marie, Kaelin, & Barsh, 2007) from POMC neurons increases body weight in female but not male mice, while deletion of GABA_B_ receptors in POMC neurons results in diet-induced obesity in male but not female mice (Ito et al., 2013). In addition, while both male and female MC4R−/− mice have increased body weight, female mice gain more weight than male mice relative to their same sex controls (Huszar et al., 1997). For each of these examples, one sex showed either no change or a reduced increase in weight rather than an opposite reduction in body weight, which is in contrast to the data presented here, where opposite changes in VTA MC3R neuron activity in males and females cause a similar decrease in feeding. Thus, the results presented here are a novel and interesting demonstration of activity and sex-dependent control of behavior by a specific population of neurons.

One potential explanation for these results is that there could be an inverted-U shaped dose-response curve for VTA MC3R neuron activity-mediated control of feeding (Figure 6), in which both males and females are on the peak of the curve, but baseline activity level is greater in females (right side of peak) than in males (left side of peak). Under this scenario, both a decrease in VTA MC3R neuron activity in females (to the left on the curve) and an increase in VTA MC3R neuron activity in males (to the right on the curve) do not alter feeding, as both remain on the peak of the curve, but an increase in neural activity in females (to the right) and a decrease in neural activity in males (to the left) drive both off of the peak of the curve to cause a decrease in feeding (Figure 6). In support of this model, both increases (Cannon, Abdallah, Tecott, During, & Palmiter, 2004; Mikhailova et al., 2016) and decreases (Szczypka et al., 1999; Zhou & Palmiter, 1995) in dopamine result in a reduction in feeding (i.e. an inverted U response), and the baseline activity of dopamine neurons is enhanced in estrus females compared to males (Calipari et al., 2017). As ~40-60% of VTA MC3R neurons are dopamine neurons (Figure 5B and (Lippert et al., 2014)), changes in dopamine release downstream of alterations in VTA MC3R neuron activity could drive these effects, although this remains to be tested.

**Figure 6:**
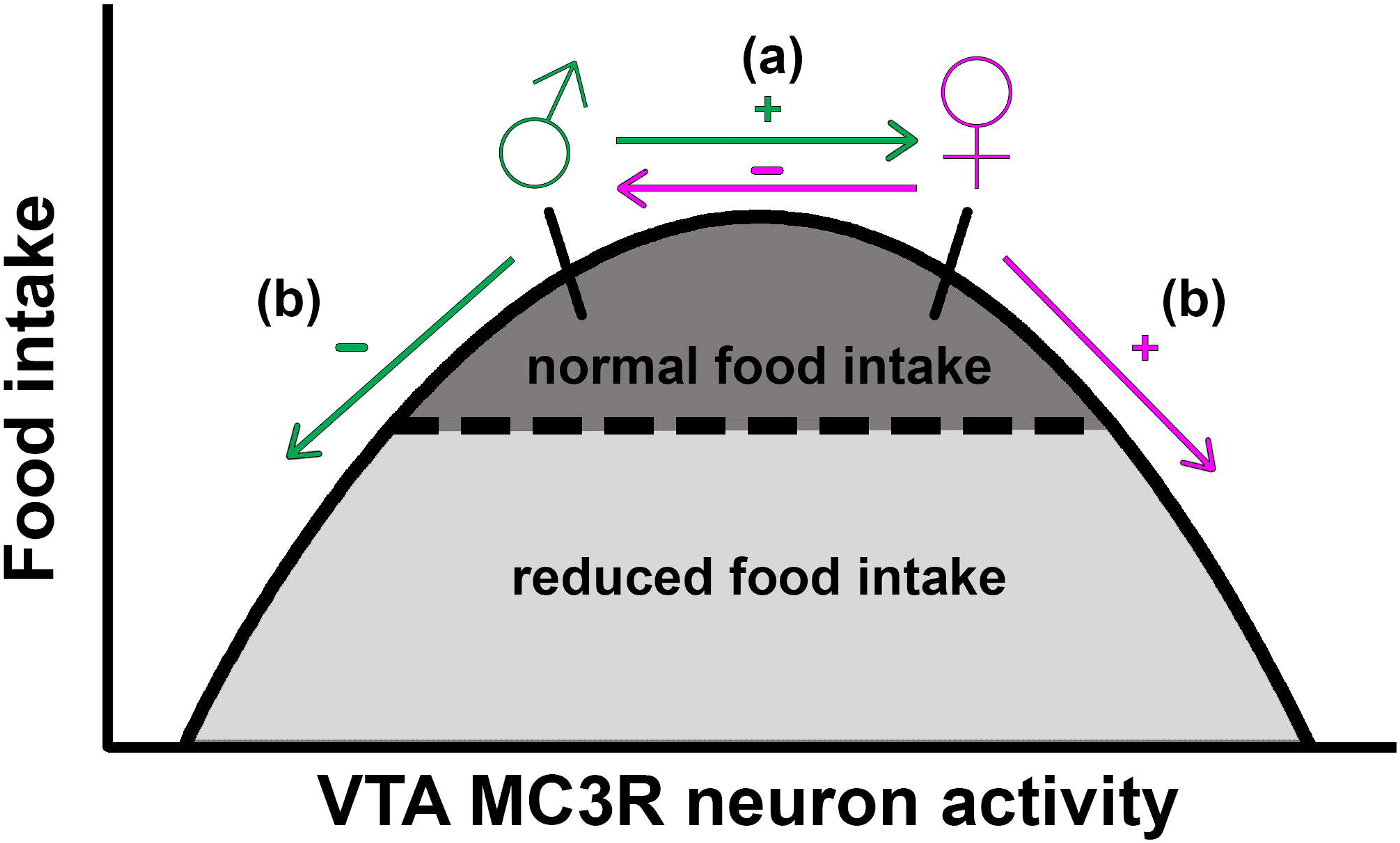
A model for sexually dimorphic VTA MC3R neuron activity-mediated control of feeding. At baseline, both males and females are at the peak of the curve which allows for normal feeding behavior, but the baseline VTA MC3R neuron activity is greater in females than males. (a) An increase in VTA MC3R neuron activity (via hm3Dq) in males (green arrow and +) and a decrease in activity (via hm4Di) in females (magenta arrow and −) would allow the mice to stay on the peak of the curve and maintain normal feeding (dark shaded area). (b) In contrast, a decrease in VTA MC3R neuron activity in males (green arrow and −) and an increase in activity in females (magenta arrow and +) would move the mice off the peak of the curve and cause a decrease in food intake (light gray shaded area).

Although the model presented above could explain these responses, the mechanisms underlying the sex differences in the acute response to VTA MC3R neuron activation are unknown, and many causes could be responsible for the sex differences in feeding observed in these studies. This includes differences in how VTA MC3R neurons respond to DREADD-mediated activation and inhibition, anatomical differences in VTA MC3R neurons or their circuit connectivity, or difference in gonadal steroid environment or the response to gonadal steroids. From our experiments, male and female VTA MC3R neurons do not appear to have differential responses to CNO treatment as we observed no differences in the ability of CNO to facilitate *c-fos* gene activation via Gq DREADD (Figure 4). Moreover, we did not detect any differences in the number of neurons expressing MC3Rs, TH, or those co-expressing both markers (Figure 5 A-C) and saw no differences in the rostro-caudal distribution of MC3R neurons (Figure 5 D), TH neurons (data not shown), or co-expressing neurons (data not shown). It also appears unlikely that differences in the circuit connectivity of VTA MC3R neurons would mediate these effects, as we have recently shown that male and female VTA MC3R neurons do not differ in their efferent projections or their afferent inputs (Dunigan et al., 2020).

One potential explanation for the differences observed here is that there could be sex differences in the specific neurotransmitters or combinations of transmitters released from VTA MC3R neurons. Approximately 40-60% of VTA MC3R neurons are dopamine neurons (Figure 5B and (Lippert et al., 2014)), and the remaining ~40-60% of VTA MC3R neurons lack TH, indicating that they are GABA and/or glutamate neurons. Alterations in VTA GABA (Soden et al., 2020; Stamatakis et al., 2013; Tan et al., 2012; van Zessen, Phillips, Budygin, & Stuber, 2012) and glutamate (Qi et al., 2016; Root, Mejias-Aponte, Qi, & Morales, 2014; H. L. Wang, Qi, Zhang, Wang, & Morales, 2015) neuron activity have been shown to differentially regulate reward- and aversion-associated behaviors. Therefore, we cannot rule out the contribution of GABA or glutamate release from VTA MC3R neurons to these effects. Additionally, many different intersectional phenotypes of VTA neurons exist including dopamine neurons that co-release glutamate (Hnasko et al., 2010; Zhang et al., 2015) or GABA (Berrios et al., 2016; Kim et al., 2015; Tritsch, Oh, Gu, & Sabatini, 2014), and neurons that co-release glutamate and GABA from the same individual neuron (D. H. Root et al., 2014). As the mechanisms controlling release of one neurotransmitter over the other and the dynamics of co-release from these neurons are not understood, it is possible that neurotransmitter release dynamics from VTA MC3R neurons differ in the two sexes and could contribute to sex differences in feeding behavior, but these possibilities will need to be examined in future experiments.

We also cannot rule out the possibility that reproductive hormones contribute to the sex differences observed here, as the mesocorticolimbic system has been shown to be highly sensitive to the effects of reproductive hormones. For example, baseline activity of dopamine neurons is enhanced during estrus (Calipari et al., 2017) and dopamine synthesis and turnover are increased (Pasqualini, Olivier, Guibert, Frain, & Leviel, 1995) and striatal DAT density is decreased (McArthur, Murray, Dhankot, Dexter, & Gillies, 2007) by estradiol. It is therefore possible that estradiol may affect the responses of VTA MC3R neurons to acute activation or inhibition, including the physiological effects facilitated by melanocortins, resulting in sexually dimorphic behavioral responses. This possibility appears unlikely, however, because the experimental design utilized in these experiments should have caused individual female mice to be in distinct stages of the estrus cycle on test days. For each individual mouse, test days were separated by 2 days and injections were counterbalanced between mice. Thus, with a ~4 day estrus cycle, the female mice used in this study should have been in distinct stages on test days, and mice were tested in 7 individual cohorts, which makes it unlikely that all female mice in each cohort would have been in the same stage of the estrus cycle during CNO test days. We did not monitor estrus cycle in these studies, however, so it is possible that reproductive hormones could have contributed to the observed differences, although this does not appear likely. Furthermore, we cannot rule out the possibility that reproductive hormone-independent, cell-autonomous sex dependent mechanisms contributed to these differential responses. Overall, further experiments will be necessary to identify the exact mechanism guiding the sex differences in the feeding response to acute activation and inhibition of VTA MC3R neurons.

In summary, we have demonstrated that VTA MC3R neurons acutely control feeding in a sex- and activity-dependent manner. Overall, these studies suggest that VTA MC3R neurons may be a key site for interaction between homeostatic and hedonic circuits controlling feeding. Furthermore, these studies also reveal important sex differences in behavior and provide novel data showing that opposing changes in neuronal activity in male and female mice can cause similar changes in behavior.

## Data Availability Statement

The data that support the findings of this study are available from the corresponding author upon reasonable request.

